# Rational Design of SARS-CoV-2 Spike Glycoproteins To Increase Immunogenicity By T Cell Epitope Engineering

**DOI:** 10.1101/2020.08.14.251496

**Authors:** Edison Ong, Xiaoqiang Huang, Robin Pearce, Yang Zhang, Yongqun He

**Author notes:** These authors contributed equally (Edison Ong, Xiaoqiang Huang). Corresponding authors: Yang Zhang; Yongqun He.

## Abstract

The current COVID-19 pandemic caused by SARS-CoV-2 has resulted in millions of confirmed cases and thousands of deaths globally. Extensive efforts and progress have been made to develop effective and safe vaccines against COVID-19. A primary target of these vaccines is the SARS-CoV-2 spike (S) protein, and many studies utilized structural vaccinology techniques to either stabilize the protein or fix the receptor-binding domain at certain states. In this study, we extended an evolutionary protein design algorithm, EvoDesign, to create thousands of stable S protein variants without perturbing the surface conformation and B cell epitopes of the S protein. We then evaluated the mutated S protein candidates based on predicted MHC-II T cell promiscuous epitopes as well as the epitopes’ similarity to human peptides. The presented strategy aims to improve the S protein’s immunogenicity and antigenicity by inducing stronger CD4 T cell response while maintaining the protein’s native structure and function. The top EvoDesign S protein candidate (Design-10705) recovered 31 out of 32 MHC-II T cell promiscuous epitopes in the native S protein, in which two epitopes were present in all seven human coronaviruses. This newly designed S protein also introduced nine new MHC-II T cell promiscuous epitopes and showed high structural similarity to its native conformation. The proposed structural vaccinology method provides an avenue to rationally design the antigen’s structure with increased immunogenicity, which could be applied to the rational design of new COVID-19 vaccine candidates.

## Introduction

The current Coronavirus Disease 2019 (COVID-19) pandemic caused by severe acute respiratory syndrome coronavirus 2 (SARS-CoV-2) has resulted in over 18 million confirmed cases and 702,642 deaths globally as of August 6 2020 according to the World Health Organization [1]. Tremendous efforts have been made to develop effective and safe vaccines against this viral infection. The Moderna mRNA-1273 induced vaccine-induced anti-SARS-CoV-2 immune responses in all 45 participants of phase I clinical trial [2], and advanced to phase III clinical trial in record time. On the other hand, the Inovio INO-4800 DNA vaccine not only showed protection from the viral infection in rhesus macaques, but was also reported to induce long-lasting memory [3]. In addition to these two vaccines, there are over a hundred COVID-19 vaccines currently in clinical trials including other types of vaccines such as the Oxford-AstraZeneca adenovirus-vectored vaccine (ChAdOx1 nCoV-19) [4], CanSino’s adenovirus type-5 (Ad5)-vectored COVID-19 vaccine [5], and Sinovac’s absorbed COVID-19 (inactivated) vaccine (ClinicalTrials.gov Identifier: NCT04456595). Among all the vaccines, a vast majority of them select the spike glycoprotein (S) as their primary target.

The SARS-CoV-2 S protein is a promising vaccine target and many clinical studies reported anti-S protein neutralizing antibodies in COVID-19 recovered patients [6]. After the SARS outbreak in 2003 [7], clinical studies reported neutralizing antibodies targeting the SARS-CoV S protein [8,9], which was selected as the target of vaccine development [10,11]. Since SARS-CoV-2 shares high sequence identity with SARS-CoV [12], it is presumed that neutralization of the SARS-CoV-2 S protein could be an important correlate of protection in COVID-19 vaccine development [13]. Many computational studies utilizing reverse vaccinology and immuno-informatics reported the S protein to be a promising vaccine antigen [14–16], and clinical studies identified anti-S protein neutralizing antibodies in COVID-19 recovered patients [17–19]. The cryo-EM structure of the S protein [20] and the neutralizing antibodies binding to the S protein [21,22] were determined. Besides neutralizing antibodies, studies have also shown the importance of CD4 T cell response in the control of SARS-CoV-2 infection and possible pre-existing immunity in healthy individuals without exposure to SARS-CoV-2 [6,23,24]. Overall, successful vaccination is likely linked to a robust and long-term humoral response to the SARS-CoV-2 S protein, which could be further enhanced by the rational structural design of the protein.

Structural vaccinology has shown successes to improve vaccine candidates’ immunogenicity through protein structural modification. The first proof-of-concept was achieved by fixing the conformation-dependent neutralization-sensitive epitopes on the fusion glycoprotein of respiratory syncytial virus [25]. A similar strategy has been applied to SARS-CoV-2 to conformationally control the S protein’s receptor-binding domain (RBD) domain between the “up” and “down” configurations to induce immunogenicity [26]. In this study, we extended structural vaccinology to rationally design the SARS-CoV-2 S protein by generating thousands of stable S protein variants without perturbing the surface conformation of the protein to maintain the same B cell epitope profile. In the meantime, mutations were introduced to the residues buried inside the S protein so that more MHC-II T cell epitopes would be added into the newly designed S protein to potentially induce a stronger immune response. Finally, we evaluated the computationally designed protein candidates and compared them to the native S protein.

## Materials and methods

### Computational redesign of SARS-CoV-2 S protein

Fig 1 illustrates the workflow for redesigning the SARS-CoV-2 S protein to improve its immunogenic potential toward vaccine design. The full-length structure model (1,273 amino acids for an S monomer) of SARS-CoV-2 S assembled by C-I-TASSER [27] was used as the template for fixed-backbone protein sequence design using EvoDesign [28]. Although the cryo-EM structure for SARS-CoV-2 S is available (PDB ID: 6VSB) [20], it contains a large number of missing residues, and therefore, the full-length C-I-TASSER model was used for S protein design instead. The C-I-TASSER model of the S protein showed a high similarity to the cryo-EM structure with a TM-score [29] of 0.87 and RMSD of 3.4 Å in the common aligned regions, indicating a good model quality. The residues in the S protein were categorized into three groups: core, surface, and intermediate [30], according to their solvent accessible surface area ratio (SASAr). Specifically, SASAr is defined as the ratio of the absolute SASA of a residue in the structure to the maximum area of the residue in the GXG state [31], where X is the residue of interest; the SASAr ratios were calculated using the ASA web-server (http://cib.cf.ocha.ac.jp/bitool/ASA/). The core and surface residues were defined as those with SASAr <5% and >25%, respectively, while the other residues were regarded as intermediate. Since the surface residues may be involved in the interactions with other proteins (e.g., the formation of the S homotrimer, S-ACE2 complex, and S-antibody interaction) and may partially constitute the B cell epitopes, these residues were excluded from design, and more rigorously, their side-chain conformations were kept constant as well. Besides, the residues that may form B cell epitopes reported by Grifoni et al. [15] were also fixed. The remaining core residues were subjected to design, allowing amino acid substitution, whereas the intermediate residues were repacked with conformation substitution. Specifically, 243, 275, and 755 residues were designed, repacked, and fixed, respectively; a list of these residue positions is shown in Supplementary Table S1.

**Fig 1.**
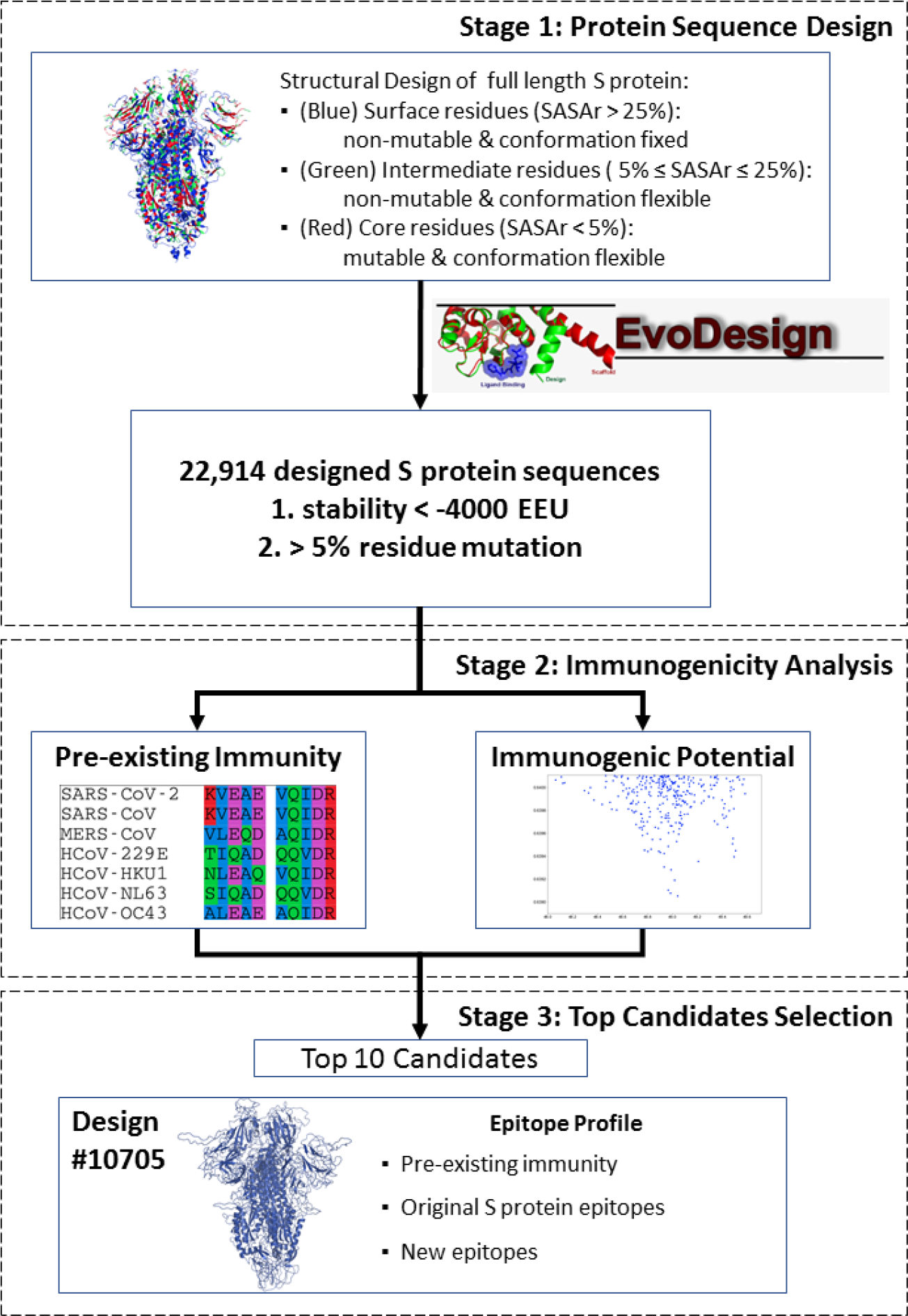
The workflow of designing and screening immunogenicity-enhanced SARS-CoV-2 S proteins. The procedure started from defining the full-length SARS-CoV-2 native S protein into surface, intermediate, and core residues. This information was then fed into EvoDesign to generate structurally stable designs that introduce mutations to the core residues while keeping the surface conformation unchanged. The output design candidates from EvoDesign were then evaluated based on their immunogenic potential. The top ten candidates were also compared and evaluated in comparison to the native S protein.

During protein design, the evolution term in EvoDesign was turned off as this term would introduce evolutionary constraints on the sequence simulation search which were not needed for this design [32]; therefore, only the physical energy function, EvoEF2 [30], was used for design scoring to broaden sequence diversity and help to identify more candidates with increased immunogenicity. We performed 20 independent design simulations and collected all the simulated sequence decoys. A total of 5,963,235 sequences were obtained, and the best-scoring sequence had stability energy of −4100.97 EvoEF2 energy unit (EEU). A set of 22,914 non-redundant sequences that were within a 100 EEU window of the lowest energy and had >5% of the design residues mutated were retained for further analysis (Fig. 1).

### MHC-II T cell epitope prediction and epitope content score calculation

The full-length S protein sequence was divided into 15-mers with 10 amino-acid overlaps. For each 15-mer, the T cell MHC-II promiscuous epitopes were predicted using NetMHCIIpan v3.2 [33], and an epitope was counted if the median percentile rank was ≤ 20.0% by binding the 15-mer to any of the seven MHC-II alleles [34] (i.e., HLA-DRB1*03:01, HLA-DRB1*07:01, HLA-DRB1*15:01, HLA-DRB3*01:01, HLA-DRB3*02:02, HLA-DRB4*01:01, and HLA-DRB5*01:01). The selection of these seven MHC-II alleles aimed to predict the dominant MHC-II T cell epitopes across different ethnicity and HLA polymorphism. The MHC-II promiscuous epitopes of the native SARS-CoV-2 S protein (QHD43416) predicted using this method were also validated and compared to the dominant T cell epitopes mapped by Grifoni et al. [15]. In brief, Grifoni et al. mapped the experimentally verified SARS-CoV T cell epitopes reported in the IEDB database to the SARS-CoV-2 S protein based on sequence homology and reported as the dominant T cell epitopes. The epitope content score (ECS) for a full-length S protein was defined as the average value of the median percentile ranks for all the 15-mers spanning the whole sequence.

### Human epitope similarity and human similarity score calculation

The human proteome included 20,353 reviewed (Swiss-Prot) human proteins downloaded from Uniprot (as of July 1, 2020) [35]. A total of 261,908 human MHC-II T cell promiscuous epitopes were predicted, as described above. The human epitope similarity between a peptide of interest (e.g., a peptide of the S protein) and a human epitope was then calculated using a normalized peptide similarity metric proposed by Frankild et al. [36]. In brief, the un-normalized peptide similarity score, *A*(*x, y*), was first determined by the BLOSUM35 matrix [37] for all the positions between a target peptide (y) and a human epitope (x), which was subsequently normalized using the minimum and maximum similarity scores for the human epitope (Eq. 1). Finally, the maximum normalized similarity score of a 15-mer peptide was calculated by comparing to all the predicted human MHC-II T cell promiscuous epitopes. The human similarity score (HSS) of the full-length S protein was calculated by averaging the human epitope similarity of all the 15-mers.

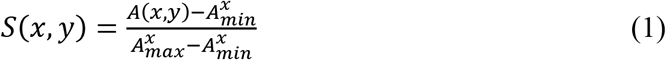

### Pre-existing immunity evaluation of the designed proteins

The pre-existing immunity of the designed proteins was evaluated and compared to that of the native S protein of seven human CoVs (i.e., SARS-CoV-2, SARS-CoV, MERS-CoV, HCoV-229E, HCoV-OC43, HCoV-NL63, and HCoV-HKU1). The sequences of the seven HCoV S proteins were downloaded from Uniprot [35] (Table S2), and the MHC-II T cell epitopes were predicted as described above. The conserved epitopes were determined by the IEDB epitope clustering tool [38] and aligned using SEAVIEW [39].

### Foldability assessment of the designed proteins

Since EvoDesign only produces a panel of mutated sequences, it is important to examine if the designed sequences can fold into the desired structure that the native S protein adopts. To examine their foldability, we used C-I-TASSER to model the structure of the designed sequences, where the structural similarity between the native and designed S proteins was assessed by TM-score [40]. Here, C-I-TASSER is a recently developed protein structure prediction program, which constructs full-length structure folds by assembling fragments threaded from the PDB, under the guidance of deep neural-network learning-based contact maps [41,42]. The ectodomain of the S homotrimers was visualized via PyMOL [43].

## Results

The epitope content score (ECS) and human similarity score (HSS) of the S proteins from seven HCoV strains (severe HCoV: SARS-CoV-2, SARS-CoV, and MERS-CoV; mild HCoV: HCoV-229, HCoV-HKU1, HCoV-NL63, and HCoV-OC43) were computed. The ECS for the severe HCoV S proteins was significantly different from that for the mild ones *(p* = 0.0016, Mann-Whitney). In terms of HSS, the severe HCoV S proteins tended to be less self-like compared to the mild ones (*p* = 0.097, Mann-Whitney). Overall, it was shown that both ECS and HSS might be used as indicators of the immunogenic potential of the designed S proteins.

On the other hand, previous studies suggested the potential role of pre-existing immunity in fighting COVID-19 [6,23,24]. Therefore, the predicted MHC-II T cell promiscuous epitopes of the SARS-CoV-2 S protein were compared to those from the other six HCoVs. There were two SARS-CoV-2 predicted MHC-II T cell promiscuous epitopes, which were also present on all of the seven HCoV S proteins (Fig 2), which could be potentially linked to pre-existing immunity. Therefore, the designs were subsequently filtered based on the availability of these pre-existing immunity-related epitopes (Fig 1). In particular, the SARS-CoV-2 promiscuous epitope S816-D830 overlapped with the dominant B cell epitope F802-E819 reported by Grifoni et al. [15].

**Fig 2.**
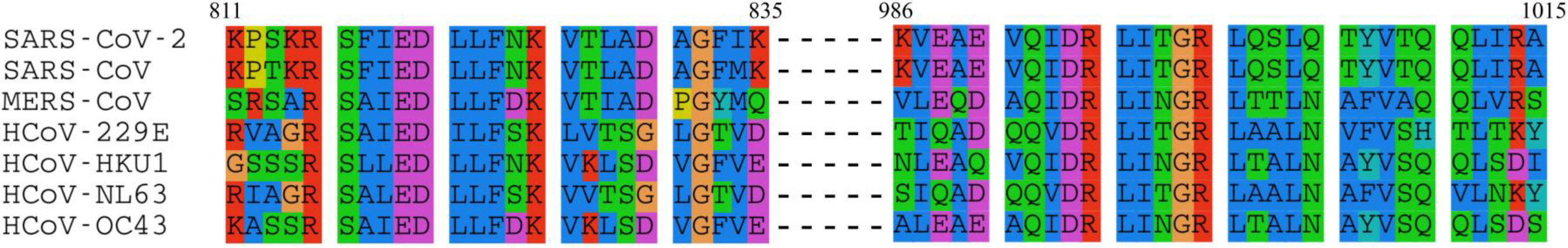
The two pre-existing immunity-related SARS-CoV-2 MHC-II T cell promiscuous epitopes. The first SARS-CoV-2 promiscuous epitope is located within residues 816-830 (indexed by SARS-CoV-2).

Among the 22,914 designs with relatively low stability energy, 19,063 candidates that contained the two pre-existing immunity-related epitopes were ranked based on ECS and HSS (Fig 3A). Using the ECS and HSS of the native SARS-CoV-2 S as the cutoff, we obtained 301 candidates with a better immunogenic potential (i.e., lower ECS and HSS) (Fig 3B). Ten candidates with balanced ECS and HSS were selected and evaluated (Table 1, full-length sequences in Table S3).

**Fig 3.**
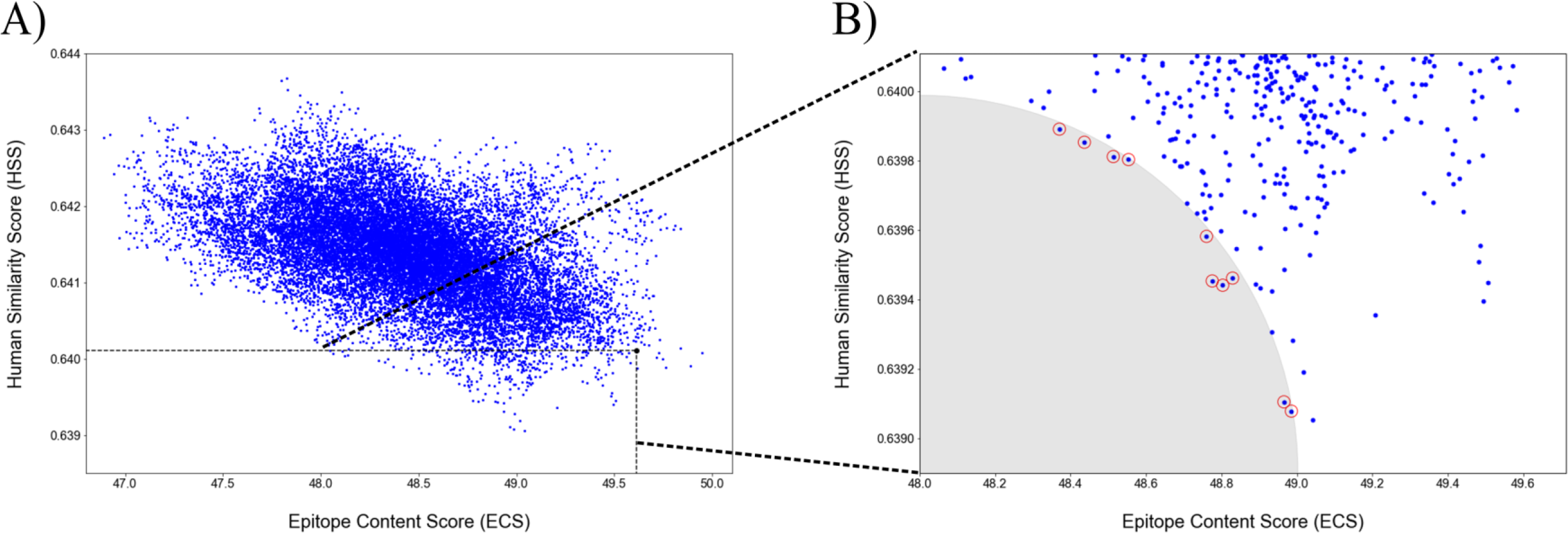
The epitope content score (ECS) and human similarity score (HSS) for designed S proteins. (A) All 22,914 designs. Each design is shown as a blue dot, whereas the native SARS-CoV-2 S was plotted as a black dot. The dashed-line box defines the 301 candidates with both lower ECS and HSS scores than the native. (B) The shaded area contains the top ten candidates with balanced ECS and HSS scores.

**Table 1.** Summary of the features for the top 10 designs. The table is ranked based on the designs’ free energy scores (from low to high) except the native S protein.

| Design ID | PEC | REC <sup>a</sup> | ECS | HSS | FE (EEU) | RMSD (Å) <sup>b</sup> | TM-score <sup>b</sup> | SI (%) |
| --- | --- | --- | --- | --- | --- | --- | --- | --- |
| 10705 | 40 | 31 | 48.78 | 0.6394 | -4051.21 | 3.45 | 0.931 | 93.9 |
| 10763 | 40 | 31 | 48.80 | 0.6394 | -4051.04 | 3.06 | 0.944 | 92 |
| 12865 | 40 | 31 | 48.76 | 0.6396 | -4044.99 | 3.14 | 0.939 | 91.9 |
| 19356 | 41 | 30 | 48.44 | 0.6399 | -4020.14 | 3.12 | 0.929 | 90.9 |
| 20348 | 38 | 30 | 48.99 | 0.6390 | -4014.74 | 3.33 | 0.929 | 94 |
| 20467 | 38 | 30 | 48.97 | 0.6391 | -4014.10 | 4.32 | 0.901 | 92 |
| 20671 | 37 | 28 | 48.83 | 0.6395 | -4013.03 | 3.36 | 0.94 | 94.7 |
| 22676 | 36 | 28 | 48.37 | 0.6399 | -4001.70 | 3.35 | 0.939 | 93.8 |
| 22769 | 38 | 28 | 48.51 | 0.6398 | -4001.11 | 3.27 | 0.937 | 93 |
| 22869 | 38 | 28 | 48.55 | 0.6398 | -4000.23 | 3.24 | 0.919 | 90.3 |
| Native | 32 | -- | 49.61 | 0.6401 | -- | -- | -- | -- |

PEC: Promiscuous Epitope Count; REC: Recovered Epitope Count; ECS: Epitope Content Score; HSS: Human Similarity Score; FE: Free Energy (EvoEF2 energy unit); RMSD: Root Mean Square Deviation; TM: TM-score; SI: Sequence identity.
<sup>a</sup>: The number of predicted promiscuous epitopes in designs that overlap with those in the native S protein.
<sup>b</sup>: The RMSD and TM-score compared to the C-I-TASSER model of the native S protein.

Design-10705 was overall the best candidate with high structural similarity to the native S protein and good immunogenic potential (in terms of promiscuous epitope count, ECS and HSS scores) amongst the top ten candidates. The candidate Design-10705 had a 93.9% sequence identity to the native S protein with TM-score (0.931) and RMSD (3.45 Å) to the C-I-TASSER model of the native S protein. The homo-trimer 3D structure of Design-10705 was visualized and compared to the S protein C-I-TASSER and cryo-EM structural models (Fig 4). In terms of immunogenicity, it had the second-highest number of promiscuous epitopes. Table 2 showed the complete MHC-II T cell epitope profile of Design-10705. There were 32 predicted promiscuous epitopes in the native S protein (Table S4), and 31 of them were recovered in Design-10705. The two pre-existing immunity-related epitopes, V991-Q1005 and S816-D830, were both recovered in the new design. Besides these two epitopes, there were 19 epitopes identical to the native S protein epitopes, while 10 epitopes had at least one mutation in Design-10705. Compared with the native S protein, the only missing MHC-II epitope in design 10705 was V911-N926, which was predicted to have reduced binding affinity to HLA-DRB1*03:01 and HLA-DRB4*01:01. Critically, this design introduced nine new MHC-II T cell promiscuous epitopes, which could potentially induce a stronger immune response with minimal perturbation compared with the native S protein.

**Fig 4.**
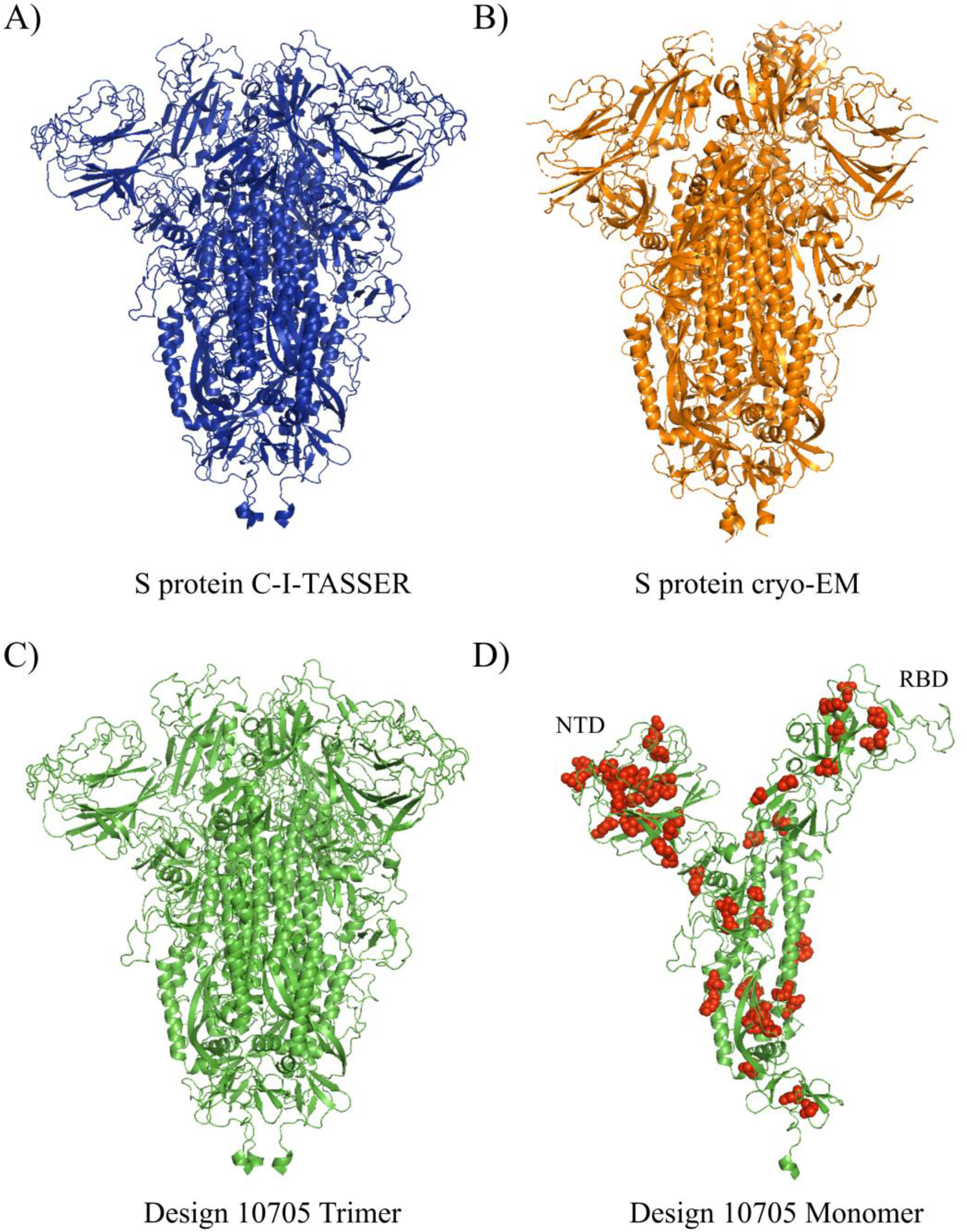
The 3D structures of A) C-I-TASSER S protein trimer, B) cryo-EM trimer, C) Design-10705 trimer, and D) Design-10705 monomer. The ectodomain of Design-10705 was modeled using C-I-TASSER. Both the homo-trimer and monomer of Design-10705 were rendered. The mutations introduced in Design-10705 are shown in red spheres.

**Table 2.**
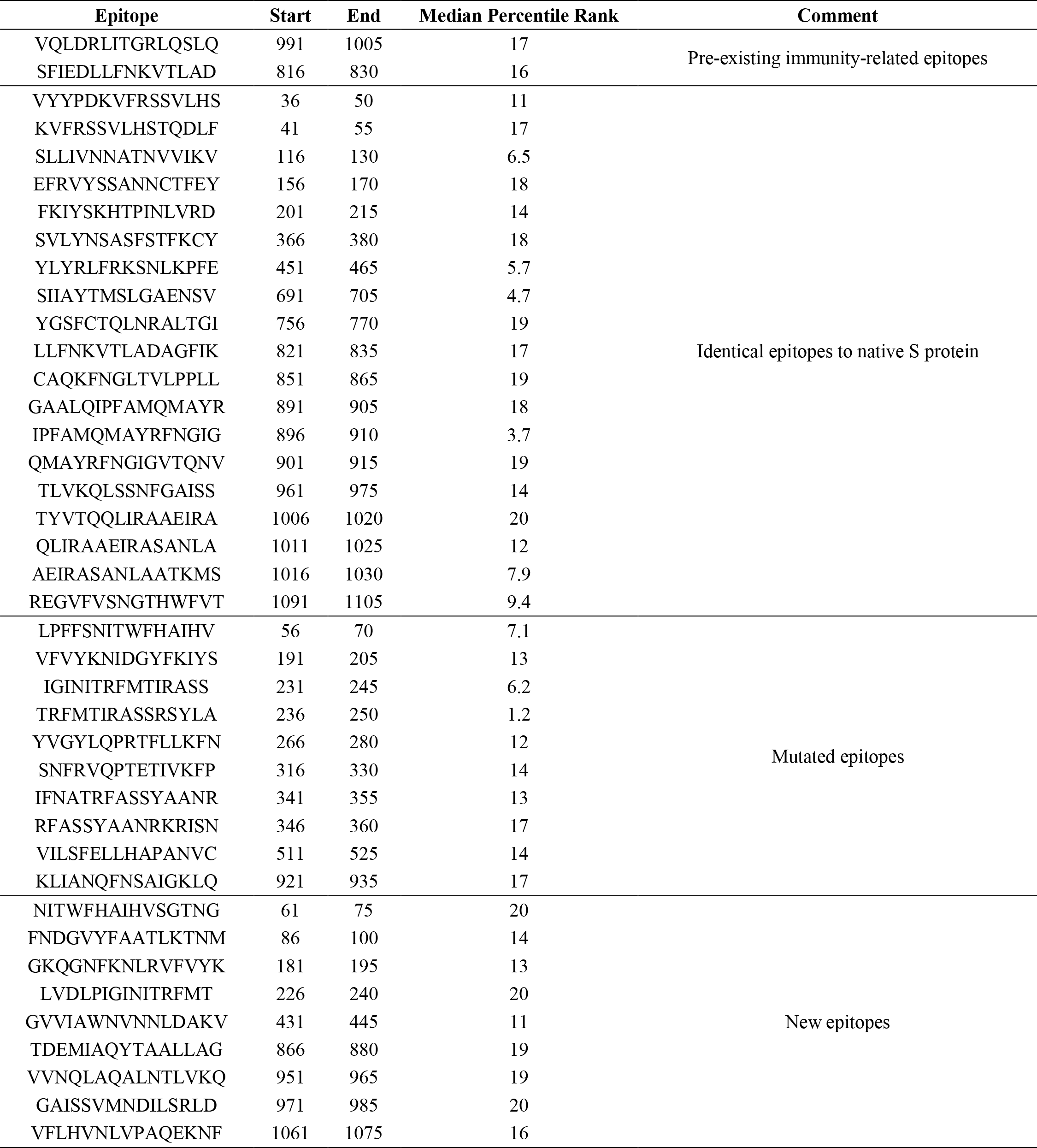
The predicted promiscuous MHC-II T cell epitopes of Design-17050.

## Discussion

The subunit, DNA, and mRNA vaccines are typically considered to be safer but often induce weaker immune responses than the live-attenuated and inactivated vaccines. Although the addition of adjuvant or better vaccination strategies can compensate for the immunogenicity, the addition of new epitopes to the antigen provides an alternative way to induce stronger immune responses [44,45]. During the protein design process, we applied design constraints so that the surface conformation, and in particular, B cell epitopes of the designed S protein variants were unchanged. For the designed S proteins with at least 5% of the core residues mutated, the immunogenicity potential of these candidates was evaluated and was structurally compared to the native S protein. The top candidate (Design-10705) recovered 31 out of 32 MHC-II promiscuous epitopes, and, the two pre-existing immunity-related epitopes (V991-Q1005 and S816-D830) were present in the design. In addition to the 31 recovered epitopes, Design-10705 also introduced nine new MHC-II promiscuous epitopes with the potential to induce stronger CD4 T cell response.

The concept of manipulating epitopes to decrease the immunogenicity has been applied to therapeutic proteins. King at el. disrupted the MHC-II T cell epitopes in GFP and *Pseudomonas* exotoxin A using the Rosetta protein design protocol [46,47]. The EpiSweep program was also applied to structurally redesign bacteriolytic enzyme lysostaphin as an anti-staphylococcal agent with reduced immunogenicity to the host [48,49]. In this study, a similar strategy, but to improve immunogenicity, was applied to redesign the SARS-CoV-2 S protein as an enhanced vaccine candidate; specifically, we aimed to increase immunogenicity by introducing more MHC-II T cell promiscuous epitopes to the protein without reducing the number of B cell epitopes.

The addition of epitopes to induce stronger immune responses has been previously applied to develop H7N9 vaccines. The H7N9 hemagglutinin (HA) vaccine elicits non-neutralizing antibody responses in clinical trials [50,51]. Rudenko et al. reported that there were fewer CD4 T cell epitopes found in H7N9 HA in comparison to the seasonal H1 and H3 HA proteins [52]. Based on this finding, Wada et al. improved the H7N9 vaccine by introducing a known H3 immunogenic epitope to the H7 HA protein without perturbing its conformation, which resulted in an over 4-fold increase of HA-binding antibody response [44]. However, the number of epitopes is not the only factor that influences the protective immunity. Studies have reported that CD8 T cell epitopes might induce regulatory T cell responses [36,53], and pathogens adapted to include CD4 and CD8 epitopes with high similarity to human peptides as a means to suppress host immunity for its survival [54]. Therefore, we examined the significance of ECS and HSS in the context of mild versus severe forms of HCoV infection and then utilized these two scores to evaluate the designed S protein candidates.

The computational design of the SARS-CoV-2 S protein could be coupled with some other structural modifications for a more rational structure-based vaccine design. The present study aims to introduce new epitopes to the S protein while keeping the surface residues unchanged to minimize the structural change of the designed proteins, and according to protein structure prediction, the designed candidates were structurally similar to the native S protein (Table 1 & Fig 4). The structural modifications performed on the native S protein, such as stabilizing the protein in its prefusion form [55], or fixing the RBD in the “up” or “down” state, could still be applied to the final candidate in this study. The combination of these structural vaccinology technologies into the current pipeline could further enhance the immunogenicity of the S protein as a vaccine target. However, a major limitation of the present study is the wet-lab experimental validation of the designed proteins. First, the newly designed protein sequences need to be folded properly with a structure comparable to that of the native S protein. Second, the capability of the newly added epitopes for binding MHC-II molecules and subsequently inducing immune responses need to be validated. Finally, these candidates should be tested for their protectiveness and safety in animal models.

Overall, this study presents a strategy to improve the immunogenicity and antigenicity of a vaccine candidate by manipulating the MHC-II T cell epitopes through computational protein design. In the current settings, the immunogenicity evaluation was carried out after the standard protein design simulations with EvoDesign. In the future, the assessment of the immunogenic potential could be incorporated into the protein design process so that the sequence decoy generated at each step will be guided by balancing both the protein stability and immunogenicity. Moreover, with proper prior knowledge of known epitopes (e.g., both MHC-I and MHC-II from the pathogen proteome), it is also possible to create a chimeric protein, which integrates epitopes from antigens other than the target protein.

## Acknowledgments

This work was supported in part by the National Institute of Allergy and Infectious Diseases (AI081062, AI134678), the National Institute of General Medical Sciences (GM136422, S10OD026825), and the National Science Foundation (IIS1901191, DBI2030790).

## Author contributions

Y.Z. and Y. H. conceived and designed the project. E.O. and X.H. performed the studies on protein sequence design and structural analyses. R.P. participated in the discussion. E.O. drafted the manuscript. All authors performed result interpretation, edited, and approved the manuscript.

## Competing financial interests

The authors declare no competing financial interests.

## Supporting Information

**S1 Table. SARS-CoV-2 S protein residues’ core, intermediate, and surface definition for EvoDesign.**

**S2 Table. Seven human coronavirus S proteins.**

**S3 Table. The full-length sequences of the top ten designs.**

**S4 Table. The predicted MHC-II T cell promiscuous epitopes of the native SARS-CoV-2 S protein.**

## Notes

### Competing Interest Statement

The authors have declared no competing interest.

